# Studying POU2F1 to Unveil the Structural Facet for Pan-Cancer Analysis Considering the Functional Annotations and Sequence-Structure Space Paradigm

**DOI:** 10.1101/806489

**Authors:** A. Dey, S. Sen, U. Maulik

## Abstract

POU domain class 2 homebox 1 (POU2F1) is largely known for its transcription factor(TF) activity. Due to its association with different types of malignancies, POU2F1 becomes one of the key factors in pan-cancer analysis. However, in spite of understanding it as a potential drug target, none of the drug has been designed till date due to its extreme dynamicity. In this article, we have proposed a three fold comprehensive framework for understanding the structural conservation and co-variation of POU2F1. Firstly, a gene regulatory network based non-pathogenic and pathogenic study have been performed to understand the strong association between cancers and POU2F1. After that, based on evolutionary sequence space study, the comparative sequential dynamicity of the protein members of POU domain family has been observed mostly between non-human and human samples. Subsequently, the reciprocity effect of the residual co-variation has been identified through direct coupling analysis. Along with that, the structure of POU2F1 has been analysed depending on quality assessment and normal mode based structure network. Comparing the sequence and structure space information, the most significant set of residue viz., 3, 9, 13, 17, 20, 21, 28, 35, 36 have been identified as structural facet for stable binding. It is observed that targeting these residues can help to prevent the monomeric aggregations. This study demonstrates-the observed malleability of POU2F1 is one of the prime reason behind its functional multiplicity in terms of protein moonlighting.

**SIGNIFICANCE:** POU2F1 is one of the important TFs associated with several malignancies. In spite of orderly stable structure in primitive family proteins, this protein is highly unstable at the monomeric stage. Previous studies show that POU2F1 has a clear association with 121 different diseases. Therefore, it becomes one of the important links in the pan-cancer study. Interestingly, TFs can be considered as a potential drug target. However, it is hardly possible to control the extreme dynamicity of TFs. In this regard, protein moonlighting plays an important role. We have tried to provide a theoretical frame to build understanding on POU2F1. The study provides pathogenic and non-pathogenic connections of the protein in terms of comprehensive and precise Gene Regulatory Network. Subsequently, the instability has been unveiled and the structural facet has been identified. Finally, the specified set of function for a suitable session to target the protein is analyzed based on protein moonlighting properties.

## INTRODUCTION

POU domain class 2 homebox 1 (POU2F1) is widely known for it transcription regulation activity. In general, functional annotations of the proteins are largely dependent on their structural orchestration. This explains how the rate of structural malleability of POU2F1 is highly responsible for its dynamic nature. This may suspected to be one of the prime reason behind the association with multiple malignancies. Few of the recent publications have discussed the importance of POU2F1 in pathogenic progression. A circular association of ECD, Akt signaling pathway and POU2F1 have been reported in Gastric Cancer (1). Functionally, phosphorylated Akt protein is behaving like Activated Cdc42 associated Kinase 1 (ACK1). Subsequently, Akt signaling pathway triggers and up-regulates POU2F1 which regulates the promoter region of ECD protein. Similarly, the combination of early high-risk gene E6/E7, P53 and POU2F1 is responsible for activating Wnt signaling pathway during cervical cancer (2). Eventually, POU2F1 becomes one of the important TFs which are regulating a large number of genes. Evidence shows that POU2F1 is associated with a total of 121 diseases such as Hepatocellular Carcinoma, Prostate Carcinoma, etc. (3). The activity of POU2F1 has been shown as a key factor during cell growth and epithelial to mesenchymal transition (EMT) during Hepatocellular Carcinoma. Similarly, the involvement of POU2F1 in colon cancer is shown in (4). The study shows that POU2F1 is highly required for gut regeneration. Moreover, microRNA mir-499a, plays an important role on liver cancer, is responsible to induce apoptosis by suppressing POU2F1 and CAPN6 (5). Due to the largely defined dynamicity and structural duality, POU2F1 can show high functional variation. In this regard, being identified as a drug target, none of the drugs can actually be designed to control its activity. Also the unusual residual distribution of this TF is not showing a defined pattern. For example, due to alanine mutagenesis at serine residue, POU2F1 loses DNA Binding capability. Mostly this mutagenesis happens at highly disorder regions and acquires local stability. Undoubtedly, this protein is one potential key to connect the pan-cancer study. Also, its strong association with multiple malignancies make it one of the potential drug targets (as mentioned before). However, TFs are known as major example of protein moonlighting activity. Protein moonlighting is a defined phenomenon of a protein with multi functionality mostly during evolutionary adaptation. Subcellular localisation of such proteins plays a vital role for detecting their moonlighting activity. In this regard, none of the study have considered specified set of functional annotations while treated it as a drug target.

For addressing the aforementioned cases, a comprehensive frame has been proposed. As discussed, we have aimed to understand the structural conservation and co-variation of this protein along with its importance as pan-cancer and protein moonlighting candidate. Firstly, the association of the protein with pathogenic annotations has been checked based on its relation in gene regulatory network (GRN, considering the sharing functional annotations) is performed an important to build the understanding about the importance of POU2F1. Subsequently, we have tried to identify the structural facets of the protein based on the evolutionary space of the POU domain family. This experiment has two folds. Firstly, we have shown an evolutionary trait of the protein based on Shannon Entropy (SE) score. After that, evolutionary highly co-varying patches have been identified using Direct Coupling Analysis (DCA) (6). Following that, the structural space of the protein has been observed based on structure network analysis. The module distribution from each. Finally, the process has been summarized based on pathway analysis (7) which can give an insight on how diverse association of the protein due to the mentioned structural malleability.

## MATERIALS AND METHODS

The proposed method is a three fold experiment where in the first fold, GRN is constructed to interpret the important and effective areas due to dys-regulation of POU2F1. In the next fold, sequence based information are analyzed considering the evolutionary space of POU domain family (Pfam id. PF00157). After that, using the Uniprot database (8), sequence of Isoform 2 (P14859) is curated and tertiary models are predicted. Among all the sequences of POU2F1, the selected sequential isoform is observed to be associated as a heteromer chain C of many known tertiary arrangements (PDB Id. 1E3O, 1GT0 etc). However, it is not the complete sequence. A sectional structural stretches has been associated with DNA binding. Finally, we have checked the probable functional multiplicity due to the dynamic nature of the POU2F1. The complete framework is discussed below in Fig 1.

**Figure 1:**
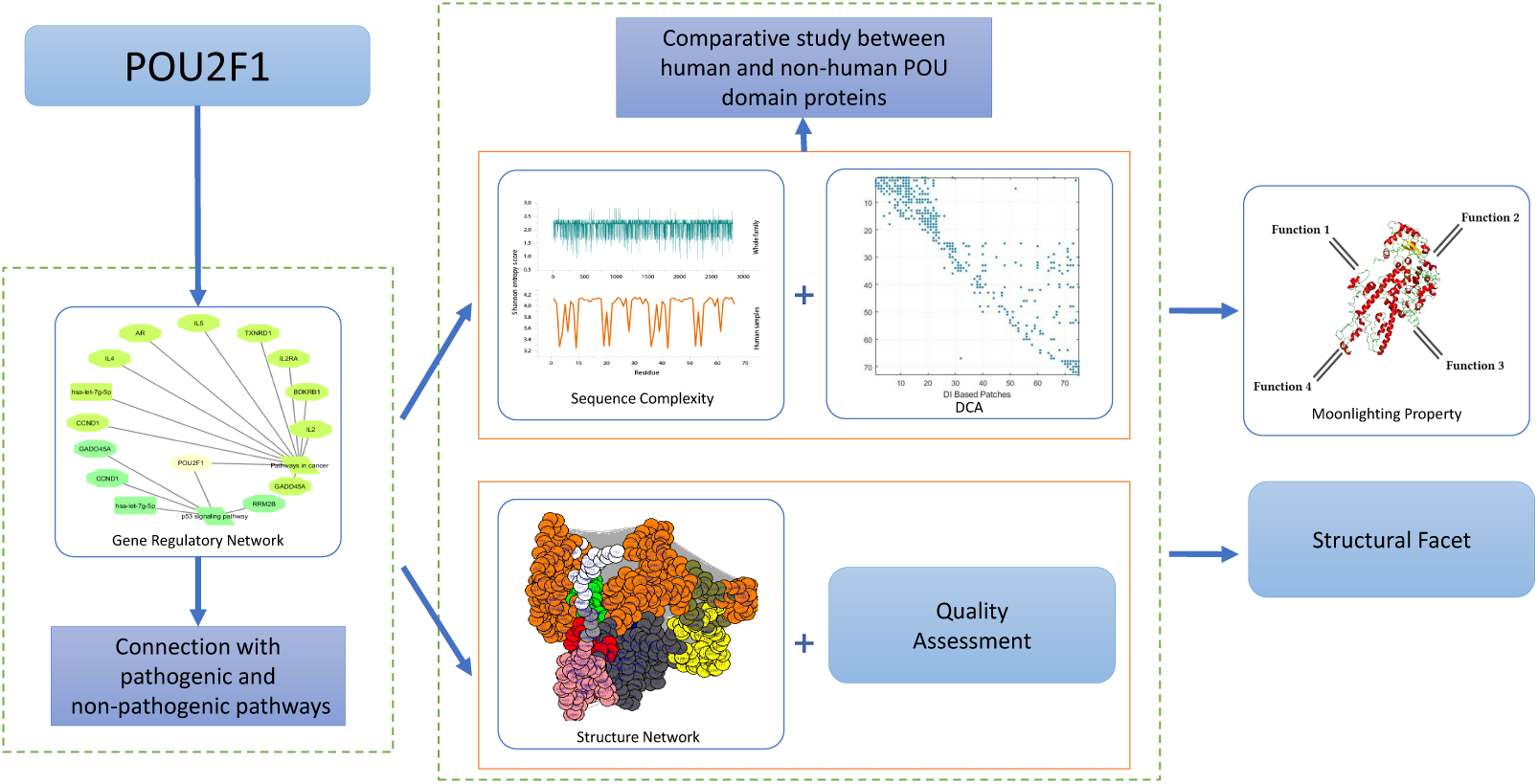
A schematic diagram to represent the flow of FunDis.

### Gene Regulatory Network

TFs are responsible to target miRNAs and genes and accordingly regulate their expression which finally lead to disease formation. To find the relation between POU2F1 and its targeted molecular regulators a GRN is constructed. In this regard, three different networks: TF-gene (9), TF-miRNA (10) and miRNA-gene (11) are performed. Based on the established networks the relation among them are identified. Moreover, these molecular regulators shared multiple biological pathways pathogenic as well as non-pathogenic. These sharing pathways may map the association between each regulator. Additionally, a final GRN is formed depending on the biological pathway information curated from KEGG pathway database (12).

### Evolutionary Trait based on Sequence Complexity

To understand the change of evolution and the associating sequence complexity of the protein through out the time period, the POU domain family sequences are curated from Pfam database (13). Some parameters are considered during the collection of the samples, such as, unaligned FASTA format is selected from the database while the sequences include tree ordering and lower case letters. It is known that Multiple Sequence Alignment (MSA) (14) and it plays an important role in a comparative functional and structural study in biological sciences. In this regard, T-Coffee alignment algorithm (15) is used to align the POU2F1 family sequences, where T-Coffee is a progressive alignment method, produces a list of pairwise alignment to conduct the MSA. Additionally, a consensus sequence (16) is obtained depending on the MSA from a Consensus Maker Tool. Customary parameters are provided to compute the consensus sequence of the family. The resulted consensus represents the logo of the protein family depending on the frequency of the amino acids. the high frequency of the amino acids are considered as the conserved region of the respective protein families (17).

Moreover, it is important to understand the structural orchestration of sequence in order to analyze the sequence present in a particular family. High entropy score signifies higher propensity toward disorderness (18). In this regard, the Shannon entropy score is calculated for the consensus sequence to unveil the trait of the family. Shannon entropy (*S*_*e*_) is defined as:

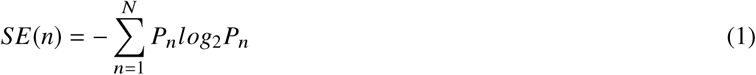

Here *P*_*n*_ is the probability and *N* is the number of the amino acids present in the sequence. The summation symbol run over the twenty amino acids that generally present in particular protein sequences. *P*_*n*_ signifies the probability of the given amino acid in the consensus sequence. Consequently, Shannon entropy score lies between 0 to *log*_2_ 20 = 4.32. It is known (19) that if the Shannon score of a sequence is more than 2.9 its propensity is more towards disorders and played an important role in the disease formation. Similarly, Shannon entropy is computed for each sequence present in the POU domain family.

### Direct Coupling Analysis

The slow changes of a protein sequence are observed throughout the evolutionary time-frame maintaining the fold of native structure unaffected (20). The unaltered amino acids are considered as conserved. These residues have an implication on sustaining the structure and function of a particular protein. Whereas, if the mutation occurs in non-conserved residues it may lead to the structural and functional disparity. The change in size, shape and other chemical properties by a mutation of a specific position is balanced by a compensatory change in another close proximity amino acid in the three-dimensional fold structure. This signifies that, in order to restore or preserve protein structure and activity, co-variation of two amino acids in evolutionary time-frame is immensely important.

To understand the dependencies of the residues and the effect of co-variation, direct coupling method is used. This is a statistical modelling to measure the strength of direct relation between two residues (21). Direct Information (DI) is used to perform the DCA. In this regard, the following equation shows how two sites of MSA is directly coupled:

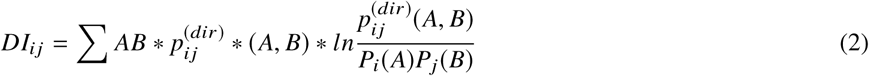

Here 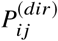 represents re-weighted frequency counts to introduce two residues for DI. *P*_*i*_ (*A*) represents the singular site frequency; i.e., probability of finding amino acid type A at *ith* position in the sequence. *P*_*j*_ (*B*) (for amino acid type B at *jth* position) is equivalent to *P*_*i*_ (*A*). *P*_*i,j*_ (*A, B*) represents joint probability of observing amino acid type A at position i and amino acid type B at *jth* position in the amino acid sequence. Coupling propensity of two residues depends on their coupling strength. Therefore, some coupled pairs are selected corresponding to top DI values. Subsequently, co-varied evolutionary patches are found and considered for further analysis. Moreover, to exhibit an effect, weighted network *G*_*DCA*_ is established considering DI score between residues. The weighted undirected network *G*_*DCA*_= (*V*_*res*_, *E*_*DI*_) where *V*_*res*_ is the set of nodes, consist of residue where *E*_*DI*_ is the weighted edges, consists of DI score between the residues. Eventually, the color modules have been formed by applying Girvan-Newman algorithm (22).

### Normal mode based Structure Network Analysis

Regarding the third fold of the study, sequential orchestration of this dynamic protein structure network is established. In this network diagram, amino acids are represented as nodes and their strength of non-covalent interactions are depicted through edges. The equation below is used to establish the interaction.

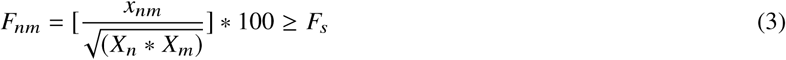

Here, *x*_*nm*_ is the number of the side chain, *n* and *m* are atom pairs of residues. *X*_*n*_ and *X*_*m*_ are the normalization factor for residues types *n* and *m* (23, 24). *F*_*s*_ is the threshold of interaction strength and 4% is the default value.

In this article, a cross-correlation matrix is calculated depending on the correlation matrix of normal mode analysis (NMA). Normal mode analysis provides a comprehensive outlook on protein tertiary structure without coarse-gaining. NMA provides some advantages considering the natural essence of the protein structure. Firstly, NMA controls Cartesian coordinates as independent variables. NMA can also represent the chain connectivity of poly-peptide chains. NMA also considers smaller number of independent variables in full atom system. Subsequently, activity of the parameters can easily be controlled tweaking dihedral bonds. Along with that, NMA considers the individual movement of each of the residues which perhaps helps to define the external and internal chemistry comprehensively. By means of the correlation matrix elastic network model is established (25), based on the tertiary structure of respective models of POU2F1 generated from I-TASSER (26). The value of cross-correlation is represented by the weight connections of particular node. Simultaneously, a full residue network is established based on the correlation network analysis. Grivan-Newman clustering method is used with threshold 0.3 to split it into densely correlated coarse-grained community cluster network.

The tertiary structure is utilized to calculate the betweenness centrality in order to identify the regions that exhibit the differences in coupled motion between the networks. Moreover, it shows the contribution to intrinsic dynamics of each node.

## RESULTS

The goal of the study is to propose a complete framework to decipher the structural conservation and co-variation of POU2F1 along with its association in Pan cancer analysis. It is evidenced that, POU2F1 is a known transcription factor, actively participates in multiple diseases such as cancer, asthama, Dermatitis, Periodontal diseases and, etc (27–29). In order to understand the nature of POU2F1, a three fold study is performed.

The involvement of POU2F1 with diverse diseases are identified from TRRUST database (9). It is found that, POU2F1 is responsible for more than 100 diseases, Fig 2 (a) represents the association of POU2F1 with other diseases by a scatter plot.

**Figure 2:**
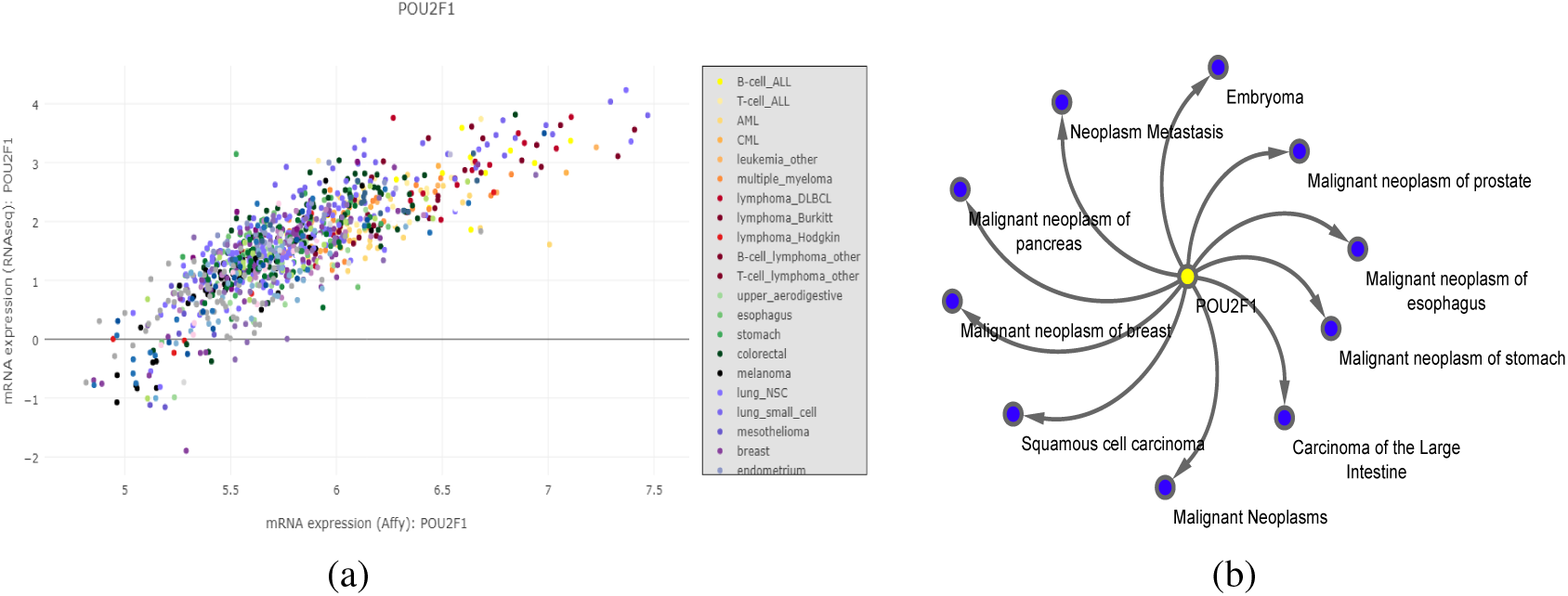
POU2F1 is associated with multiple diseases. (a) Associations are represented through a scatter plot and (b) Top ten associated diseases are shown through a network diagram.

Among the diverse diseases, top twenty diseases based on their adjusted p-value are shown by a network in Fig 2 (b) and the details are reported in Supplementary 1.

In order to interpret the regulators through which this TF is regulating the biological pathways that finally leads to disease formation an gene regulatory network is established (30). In this regard, the targeted miRNAs and genes of POU2F1 are identified. As a result 11 miRNAs and 35 genes are found, shown in Fig 3 (a). Here, POU2F1 is represented with a yellow color, miRNAs and genes are represented with green and orange color respectively. Based on this network, another network is established between selected miRNAs and genes. From the three relations (TF-gene, TF-miRNA and gene-miRNA) it is found that, POU2F1 is directly as well as indirectly regulated CCND1 genes through miRNA hsa-let-7g-5p. The network is shown in Fig 3 (b).

**Figure 3:**
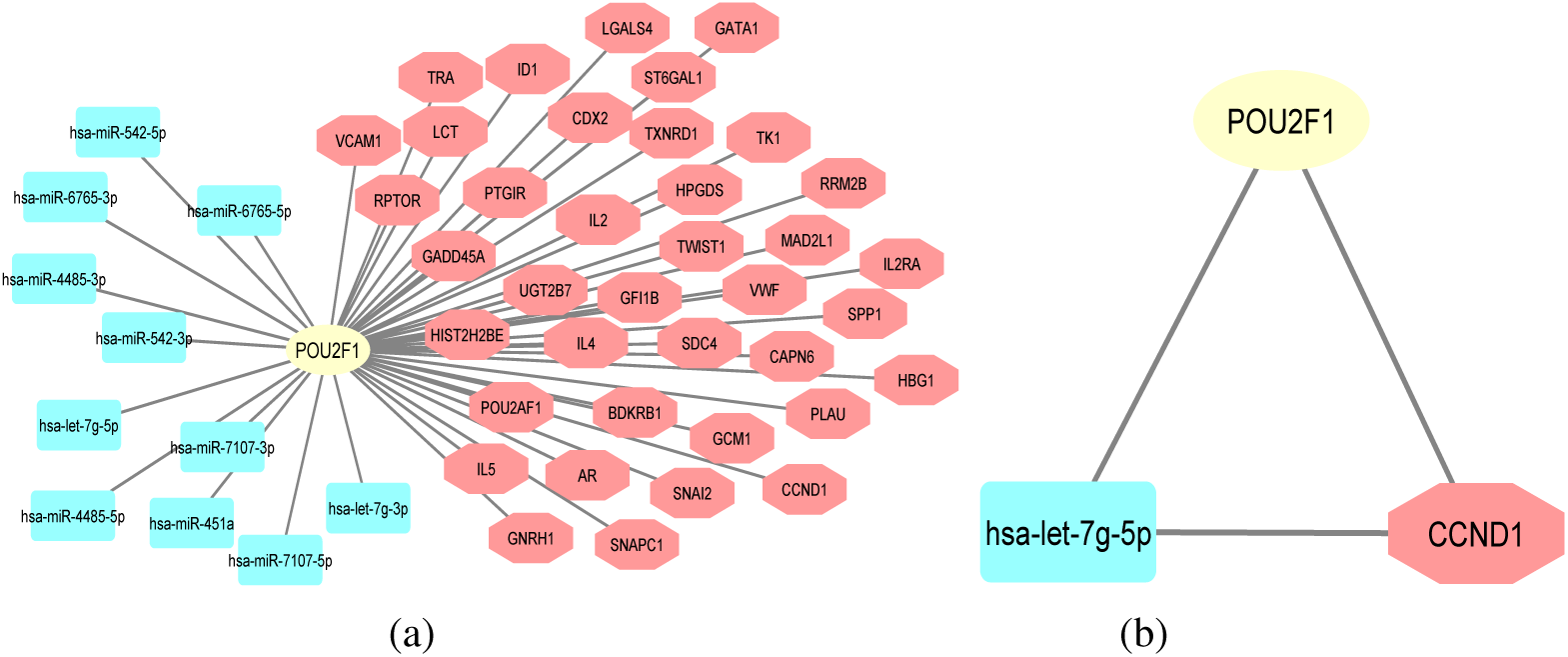
TF (POU2F1) are responsible to target miRNAs and genes (a) the targeted genes and miRNAs are represented with red and green respectively, (b) the regulatory network depicts the direct and indirect (through miRNA) regulation of gene by POU2F1

Besides constructing a regulatory network based on the literature and experimental evidences, the biological pathways are also considered for each molecular regulators. Each regulator is associated with multiple pathogenic as well as non-pathogenic pathways. Interestingly, 16 pathways are found shared among them. From these sharing pathways the final GRN is established which infer that not only one gene or miRNA are responsible for a regulatory network but also multiple relations are present those are unknown still now. Top five pathways are reported in Fig 4 based on the number of gene and miRNA association (details are reported in Supplementary 2). Each color represents a particular pathway and its associated regulators.

**Figure 4:**
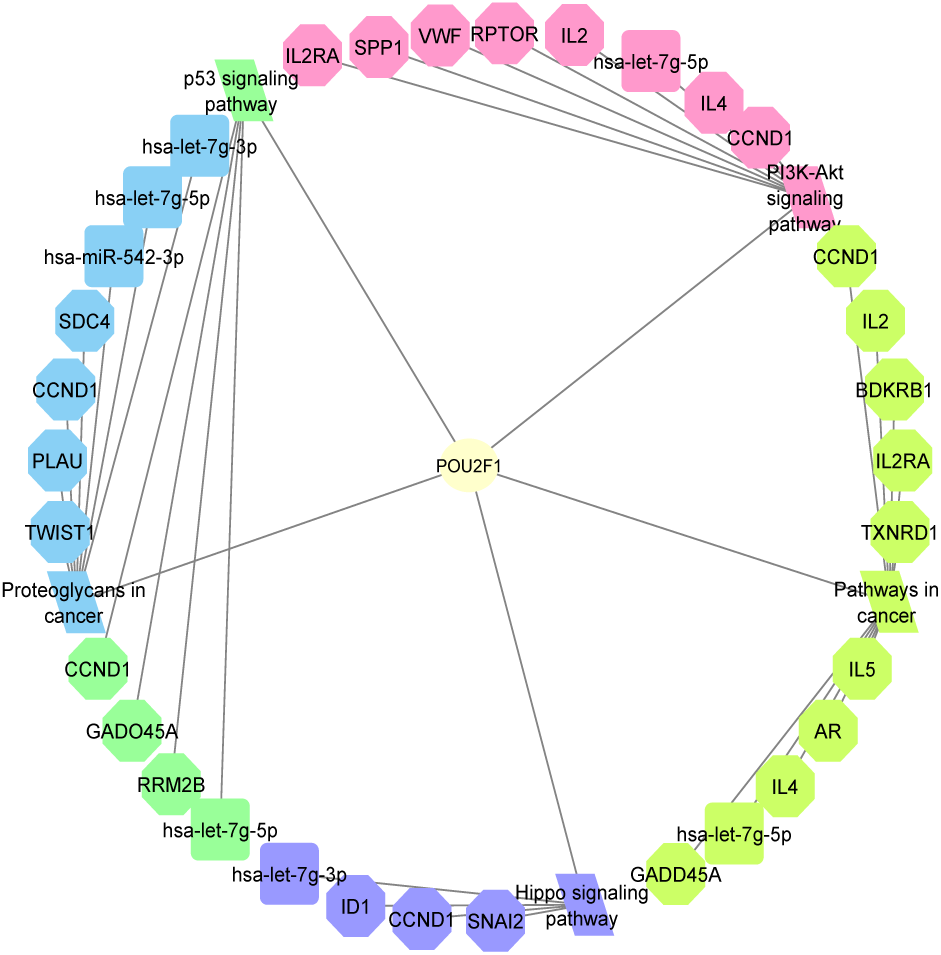
The network representation of the top five common functional pathways among POU2F1 and its targeted miRNAs and genes. Different colors represent diverse pathways and their associations.

Moreover, evolutionary trait of the POU domain family is analyzed by calculating the Shannon entropy score. During the SE calculation the human and non-human samples are divided into two groups to understand the propensity of order and disorderness of human and other primitive spices separately. The entropic scores are reported in Supplementary 3 and Fig 5 depicts the change of entropy scores of group one and two by blue and black lines respectively. In the next phase of the sequence spaced study, the co-variation propensity of residue couples at a specific evolutionary conserved position has been calculated by using DCA. From the calculation, the Direct Information (DI) scores are generated for possible coupling pairs. Among them, 93 coupled pairs are selected as highly evolutionary co-varying patches in terms of DI scores. The detail score of DCA calculation is reported in Supplementary 4 and a graph is generated for the visual representation shown in Fig 5 (b). In parallel, IUPred (31) and PONDR (32) is used to predict the disorderness of the residues present in the sequence of POU2F1 (reported in Supplementary 5). The overlapping residues from both the prediction model is considered. These common residues are clustered depending on their DCA score. The residues in same cluster represent same rate of evolution shown in Fig 6.

**Figure 5:**
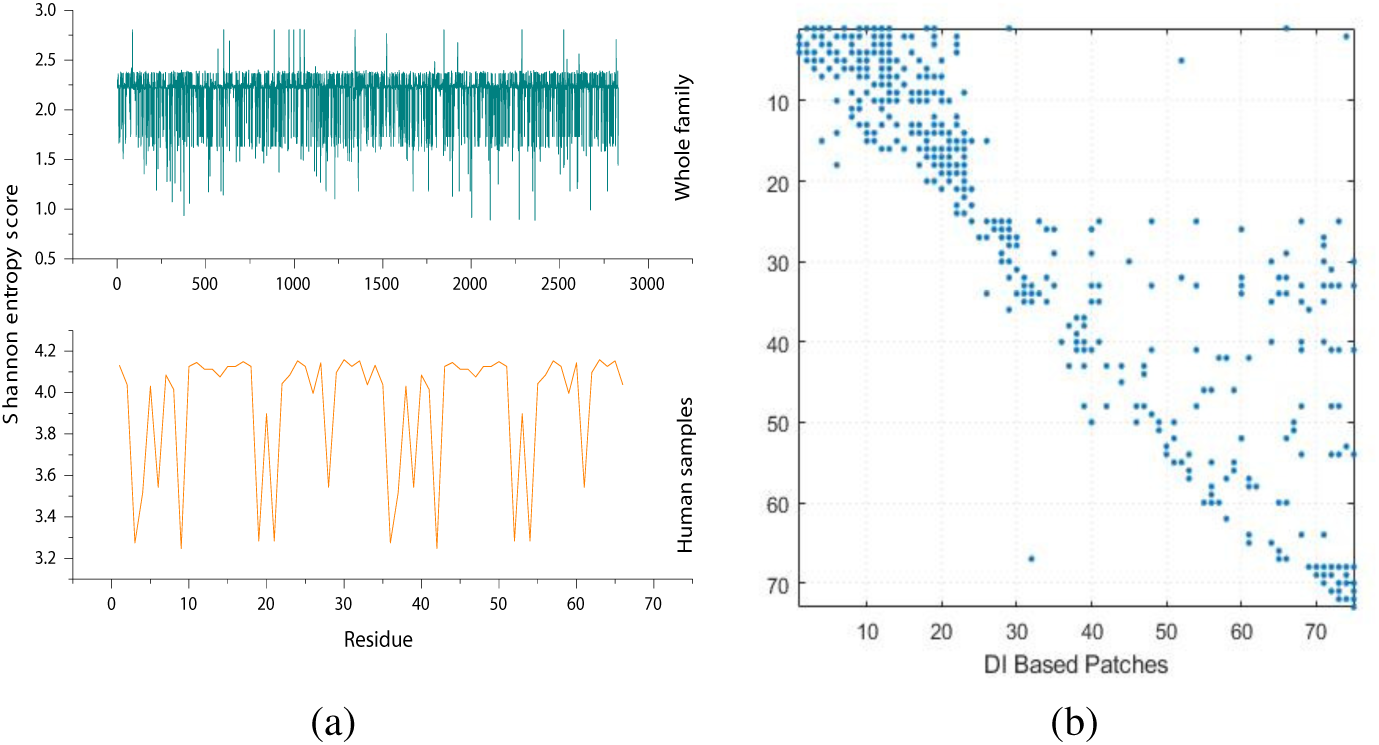
Representing the change in evolutionary trend by (a) The Shannon entropy score of Homo Sapiens samples and all other species by blue and black lines respectively and (b) The DCA score of POU2F1.

**Figure 6:**
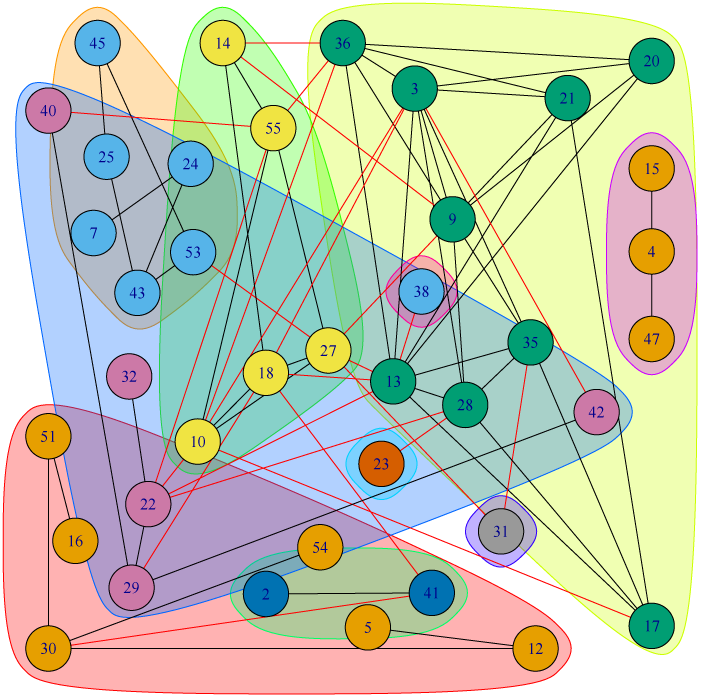
A weighted network *G*_*DCA*_ and corresponding color modules based on over all residual co-variation from DI score.

In parallel, to decipher the residual organization and dependencies at different secondary structural orchestration, structure network analysis is employed. The PDB structure of POU2F1 is established using I-TASSER (33, 34). Top five structures are constructed and depending on the C-score of the models, model 1 is selected, shown in Fig 7 (a). The green, red and yellow color of the figure represent the loop, helix and beta sheets respectively. Other models are shown in Supplementary 6 with their C-score. Accordingly, the structure network is performed to map the sequence space changes in structure space. The structure network along with the betweenness centrality graph of model 1 are shown in Fig 7 (a) and (b). Similarly, the analysis is performed for other models too, those are reported in Supplementary 7.

**Figure 7:**
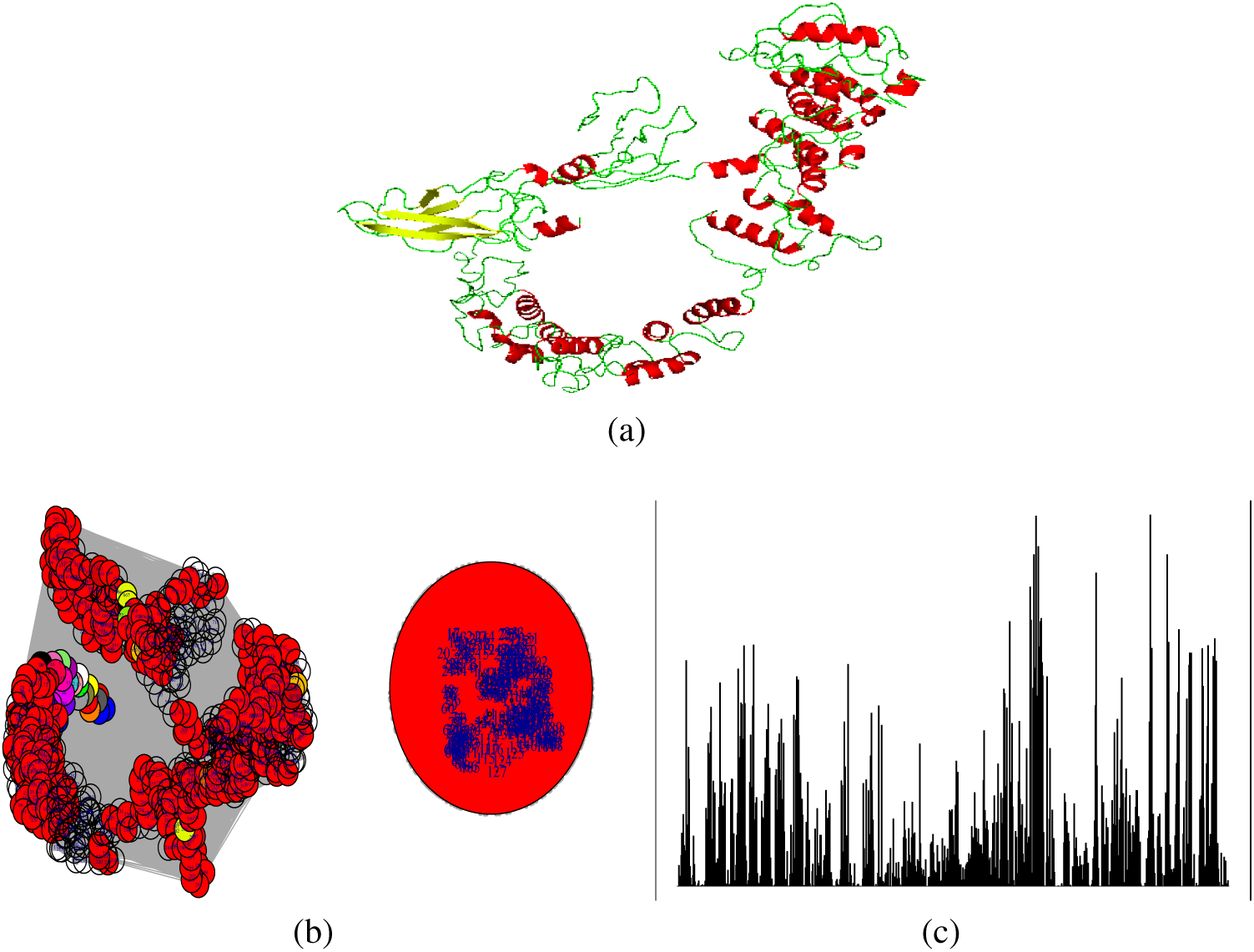
(a) The structure model 1 of POU2F1 generated from I-TASSER, where green, red and yellow color represent lop, helix and sheet respectively, (b) Structure Network of the protein and (b) betweenness centrality plot of the high confidence model performed from the database.

From the outcome, each directly coupled pair, obtained from structure space analysis, are found to be situated either in common cluster or in two densely connected clusters. The betweenness centrality is calculated to unveil the influence of a particular node on the internal dynamics of different POU2F1 structures. As the residues are aggregated into one module which is depicted from the structure network, so it is difficult to understand the structural facet of this protein. In this regard, quality assessment is performed of the PDB structure. In Supplementary 8, the courses of van der Waal forces are reported of the residues common in both the prediction models of disorderness. This forces are analyzed based on the clusters discussed in Fig 6. Finally, the structural facet is recognized depending on the cluster distribution of the residues and the corresponding non-covalent forces.

No evidence is reported till now regarding the moonlighting property of POU2F1. Moreover, from the aforementioned results, it is clear that POU2F1 is responsible to regulate diverse biological pathways, those are related to multiple diseases. Strikingly, POU2F1 is found in nucleoplasm only (35). Due to this, a BLAST is performed and based on the local similarity two proteins are found in the database having moonlighting property (36). Those proteins along with their human specific activities (function 1 (F1) and function 2 (F2)) are reported in Table 1.

From the above analysis, we can conclude that as POU2F1 is a known TF for Pan-cancer, it is associated with diverse functional pathway. During analyzing the associations, multiple targeted molecules are identified as common between pathways. POU2F1 regulated the expression of genes directly as well as indirectly and also responsible for the impact of targeted miRNAs. These miRNAs are involved in multiple diseases, the details are reported in Supplementary 9. As their expression is controlled by POU2F1 so it is inferred that POU2F1 is playing an important role in the formation of those reported diseases. If POU2F1 is targeted then it might not only control the direct formed diseases, also able to control diseases responsible for the selected miRNAs shown in Fig 3 (a).

## DISCUSSION

POU2F1 is identified as a common TF in a pan-cancer study. In Fig 2, POU2F1 and associated list of cancer is reported. Vazquez-Arreguin et al. shows the importance of POU2F1 for regeneration of colon malignancy (37). In that study, it has been shown that multiple genes have been studied based on POU2F1 regulation and its affects in pan-cancer analysis. In initial study, the pathogenic association between 116 diseases and POU2F1 has been observed. Among them, around 18 diseases are well known malignancies. The complete list is shown in Supplementary 1. Therefore, the pathogenic importance of POU2F1 is quite clearer. Although non-pathogenic association of the sample is not widely explored in any study. However, the disease causing dys-regulation are influenced through non-pathogenic pathway disruptions. In this regard, the non-pathogenic pathway association has been observed based on its transcription activities. Also specified list of malignancy oriented pathways has been revised based on mentioned GRN. The GRN associated importance of POU2F1 has been shown considering the sharing functional annotation of miRNA, gene and POU2F1 (as TF). In this regard, total 12 non-pathogenic pathways have been identified (shown in Supplementary 2). However, four pathogenic pathways are also identified. Filtering is performed to establish the strong amalgamation between POU2F1 and GRN based on the non-pathogenic connectivity. In (Fig 3 and Fig 4), functional annotation and sequential development of the GRN has been shown.

One of the objectives of the study is to understand the functional dynamic nature of the target protein POU2F1 and identify the structural facet accordingly. Initially, Functional dynamicity is analyzed based on the structural orchestration and the environmental parameters like subcellular localization, cellular compartmentalization, and so on. The residual distributions of the proteins are highly family dependent which can also consider as an evolutionary mark of that family. In (18), it is shown that, SE score of the sequences helps to understand the evolutionary trait of the protein family. It consists of the entire family specific sequence space of the respective family. Following that idea, SE is applied to each member of the POU domain family (PFAM Id.-PF00157). It is observed that POU domain samples at homo sapiens are showing an different trait than other primitive species (shown in Fig 1a). It is observed that human samples of POU domain family are having more propensity towards disorder than other primitive species. It provides insights on structural variation based on the density of the hydrophobic and hydrophilic residues. Sequence space information based on SE score is not enough to infer the internal dynamics of the protein. For this purpose, DCA has been performed. This coupling analysis shows the internal coupling propensity of the residues. Basically, the rate of co-variation has been observed using DCA disentangling coupling combinations of each residue. These calculations considered the multiple sequence alignment of the family-specific POU domain sequences. Top scoring coupling combinations are considered as highly co-varying residues and patches are shown in Fig 1b. These co-varying residue patches are mostly responsible for the structural facets. The average sequence length of POU domain proteins is 650, however, the evolutionary structural facets are restricted within a small stretch of 1 to 70 residues. The sequential abundances in terms of evolutionary co-variation are easy to interpret from the patches.However, sequence space outcome of the protein family is interconnected. More elaborately, the extended affects of the covarying residues can be observed from the clustered modules of G_*DCA*_. From the graph, there are nine specified strongly inter-connected modules has been observed. Therefore, modulating any of the residue from each of modules would be notified as cause of modulating effect to rest of the members from the same modules.

From the sequence space, we got some sets of residues assuming there evolutionary conserved comprehensive effects during folding strategy. However, this unidimensional information is incomplete. This sequence space has to be mapped on structural space to infer the structural space. POU2F1 has no proper tertiary structure, therefore among the predicted structural models from I-TASSER, the structure with higher c-score has been considered. It is observed that, the PDB structure of the selected model has higher number of flexible loops than rest of the structure. Interestingly, the covarying regions from the sequence space mostly belong to loops. Subsequently, a quality assessment analysis makes it clear that which of the residues have higher propensity to lead a healthy binding. In quality assessment, a dihedral chi1 angle clash has been found at residue number 20 which is directly associated with a sequence space module with 17, 28, 13, 35, 9, 3, 36, 21, 20 residues. Eventually based on inner coupling strength from sequence space, the inner allosteric effects may carry forward though out all the residues. In Fig 7, the respective structure network is given, considering the residual oscillation in terms of normal mode. Additionally, betweenness centrality dependent module detection has been performed. All the residues are distributed in 254 modules. Among them, only module 3 has a maximum number of residues. Apart from few residues, most of the them even the beta sheet are conserved in one module. Residues selected based on coupling propensity have also belonged to the same module. From the detailed two-step analysis, it is inferred that POU2F1 is extremely vulnerable, both structurally and evolutionary wise. Therefore, the structure of the protein has a tendency to form aggregated globule structure to achieve global stability at monomeric stage. However, comparing the sequence space and structural space information, the structural facet can be considered as residue which are shown at the initial level is of the structural study. Mapping such information at structure network, intra-molecular allosteric is highly dependent on the residues selected earlier. Also those three conserved modules atleast have one of the residues as member. Considering the fact, the known DNA binding domain of POU2F1 is at residue range 370-438. Therefore, targeting the proposed residue range cannot affect the DNA binding domain directly.

Such structural flexibility may cause of functional duality. TFs are well known for their protein moonlighting property. Considering the generic fact, POU2F1 has higher propensity for having protein moonlighting propensity. To understand whether this protein is moonlighting protein or not, we have checked it with moonPROT. As per expectation, POU2F1 has been identified with the protein moonlighting property through sequence similarity.In Table 1, probable functions of the POU2F1 has been given. In moonPROT, POU2F1 is sharing the significant sequential similarity with two proteins i.e., ATF2 and CDC1. Interestingly, localized at nucleus both the proteins have transcription and non-transcription activities. For ATF2, the switching activity of ATF2 has shown in terms of recruiting MRE11 protein fro repairing DNA damage (38). Therefore, the sequence similarity has enlightened the structural traits lies in the non-DNA binding zone of the POU2F1. Therefore the moonlighting property can shows a specific session when this proteins can be targeted.

## CONCLUSION

In this article, we have studied POU2F1 elaborately. Initially, the pathogenic and non-pathogenic connection with the protein has established based on GRN and respective sharing functional annotations in malignancies. This shows that the POU2F1 plays a vital role in a pan-cancer analysis which also makes it an important drug target. Subsequently, the evolutionary sequence-structure space analysis has provided a structural facet (a set residues) which can considered a strong binding site. Targeting the structural facet can help to prevent or slow down the monomeric aggregation rate. During experiment, the structural instability of monomeric POU2F1 is unveiled. Through the protein moonlighting property of POU2F1, it can be revealed the significant functional session when this protein supposed to be targeted.

## AUTHOR CONTRIBUTIONS

Author2 designed the research. Author2 and Author1 carried out all the experiment. Author1 and Author2 wrote the article. Author3 supervised and corrected the manuscript

## ACKNOWLEDGMENTS

We would like to acknowledge DST-INSPIRE fellowship to support the work of Ashmita Dey and Sagnik Sen.

## SUPPLEMENTARY MATERIAL

**Supplementary 1. Diverse diseases associated with POU2F1.**

**Supplementary 2. The common sharing pathways of POU2F1 and its targeted miRNAs and genes.**

**Supplementary 3. Shannon entropy score of the POU domain family proteins to understand the change in evolutionary trait.**

**Supplementary 4. The direct coupling analysis score of the residue to determine their coupling propensity.**

**Supplementary 5. The disorder score of the residues from IUPred and PONDR databases.**

**Supplementary 6. The top predicting models of POU2F1 from I-Tasser with their corresponding confidence score.**

**Supplementary 7. The structure network and betwenness centrality graphs of other POU2F1 structure models.**

**Supplementary 8. The non-covalent van der Waal forces of the residues.**

**Supplementary 9. List of diseases due to the selected 11 miRNAs.**

## REFERENCES

1. Xu, S. H., J. Z. Huang, M. L. Xu, G. Yu, X. F. Yin, D. Chen, and G. R. Yan, 2015. ACK1 promotes gastric cancer epithelial–mesenchymal transition and metastasis through AKT–POU2F1–ECD signalling. The Journal of Pathology 236:175–185.

2. Zhang, R., H. Lu, Y. Y. Lyu, X. M. Yang, L. Y. Zhu, G. G. Yang, P. C. Jiang, Y. Re, W. W. Song, C. C. Z. J. H. Wang, F. Gu, T. J. Luo, Z. Y. Wu, and C. J. Xu, 2017. E6/E7-P53-POU2F1-CTHRC1 axis promotes cervical cancer metastasis and activates Wnt/PCP pathway. Scientific Reports 7:1–13.

3. Zhong, Y., H. Huang, M. Chen, J. Huang, Q. Wu, G. R. Yan, and Chen, 2017. POU2F1 over-expression correlates with poor prognoses and promotes cell growth and epithelial-to-mesenchymal transition in hepatocellular carcinoma. Oncotarget 8:440820–44095.

4. Vazquez-Arreguin, K., C. Bensard, J. C. Schell, E. Swanson, X. Chen, J. Rutter, and D. Tantin, 2018. Oct1/Pou2f1 is selectively required for gut regeneration and regulates gut malignancy. bioRxiv 1–24.

5. Liu, Y., X. S. Y. Wang, C. Mei, L. Wang, Z. Li, and X. Zha, 2016. miR-449a promotes liver cancer cell apoptosis by downregulation of Calpain 6 and POU2F1. Oncotarget 7:13491–13501.

6. Morcos, F., T. Hwa, J. N. Onuchic, and M. Weigt, 2016. Direct coupling analysis for protein contact prediction. Nucleic Acids Research 44:W375–W382.

7. Pankratova, E. V., A. G. Stepchenko, T. Portseva, V. A. Mogila, and S. G. Georgieva, 2016. Different N-terminal isoforms of Oct-1 control expression of distinct sets of genes and their high levels in Namalwa Burkitt’s lymphoma cells affect a wide range of cellular processes. Nucleic Acids Research 44:9218–9230.

8. Consortium, T. U., 2008. The Universal Protein Resource (UniProt). Nucleic Acids Research 36:190–195.

9. Han, H., J. W. Cho, S. Lee, A. Yun, H. Kim, D. Bae, S. Yang, C. Y. Kim, M. Lee, E. Kim, S. Lee, B. Kang, D. Jeong, Y. Kim, H. N. Jeon, H. Jung, S. Nam, M. Chung, J. H. Kim, and I. Lee, 2018. TRRUST v2: an expanded reference database of human and mouse transcriptional regulatory interactions. Nucleic Acids Research 41:D380–D386.

10. Tong, Z., Q. Cui, J. Wang, and Y. Zhou, 2019. TransmiR v2.0: an updated transcription factor-microRNA regulation database. Nucleic Acids Research 47:D253–D258.

11. Ch, C. H. C., and et al., 2018. miRTarBase update 2018: a resource for experimentally validated microRNA-target interactions. Nucleic Acids Research 46:D296–D302.

12. Kanehisaa, M., and S. Goto, 2000. KEGG: Kyoto Encyclopedia of Genes and Genomes. Nucleic Acids Research 28:27–30.

13. Gebali, S. E., and et al., 2019. The Pfam protein families database in 2019. Nucleic Acids Research 47:D427–D432.

14. Chatzou, M., C. Magis, J. M. Chang, C. Kemena, G. Bussotti, I. Erb, and C. Notredame, 2016. Multiple sequence alignment modeling: methods and applications. Briefings in Bioinformatics 17:1009–1023.

15. Notredame, C., D. G. Higgins, and J. Heringa, 2000. T-Coffee: A Novel Method for Fast and Accurate Multiple Sequence Alignment. Journal of Molecular Biology 302:205–217.

16. Schneider, T. D., 2002. Consensus Sequence Zen. Applied Bioinformatics 1:111–119.

17. Dogan, T., and B. Karacali, 2000. Automatic Identification of Highly Conserved Family Regions and Relationships in Genome Wide Datasets Including Remote Protein Sequences. Proteins 42:38–48.

18. Sen, S., A. Dey, S. Chowdhury, U. Maulik, and K. Chattopadhyay, 2019. Understanding the evolutionary trend of intrinsically structural disorders in cancer relevant proteins as probed by Shannon entropy scoring and structure network analysis. BMC Bioinformatics 19:231–242.

19. Romero, P., Z. Obradovic, X. Li, E. Garner, C. Brown, and A. Dunker, 2000. Sequence complexity of disordered protein. Proteins 42:38–48.

20. Maity, H., and et al., 2005. Protein folding: The stepwise assembly of foldon units. Proceedings of the National Academy of Sciences of the United States of America 4741–4746.

21. Morcos, F., A. Pagnani, B. Lunt, A. Bertolino, D. S. Marks, C. Sander, R. Zecchina, J. N. Onuchic, T. Hwa, and M. Weigt, 2011. Direct-coupling analysis of residue coevolution captures native contacts across many protein families. Proceedings of the National Academy of Sciences of the United States of America 108:1293–1301.

22. Girvan, M., and M. E. J. Newman, 2002. Community structure in social and biological networks. Proc Natl Acad Sci U S A 99:7821–7826.

23. Brinda, K. V., and S. Vishveshwara, 2005. A network representation of protein structures: implications for protein stability. Biophysical Journal 89:4159–4170.

24. Kannan, N., and S. Vishveshwara, 1999. Identification of side-chain clusters in protein structures by a graph spectral method. Journal of Molecular Biology 292:441–464.

25. Chakrabarty, B., and N. Parekh, 2016. NAPS: Network Analysis of Protein Structures. Nucleic Acids Research 44:W375–W382.

26. Roy, A., A. Kucukural, and Y. Zhang, 2010. I-TASSER: a unified platform for automated protein structure and function prediction. Nature Protocols 5:725–738.

27. Sharpe, D. J., K. S. Orr, M. Moran, S. J. White, S. McQuaid, T. R. Lappin, A. Thompson, and J. A. James, 2014. POU2F1 activity regulates HOXD10 and HOXD11 promoting a proliferative and invasive phenotype in Head and Neck cancer. Oncotarget 5:8803–8815.

28. Holt, R. J., Y. Zhang, A. Bini, A. L. Dixon, C. Vandiedonck, W. O. Cookson, J. C. Knight, and M. F. Moffatt, 2011. Allele-specific transcription of the asthma-associated PHD finger protein 11 gene (PHF11) modulated by octamer-binding transcription factor 1 (Oct-1). Journal of Allergy Clinical Immunology 127:1054–1062.

29. Andersen, B., and M. G. Rosenfeld, 2001. POU Domain Factors in the Neuroendocrine System: Lessons from Developmental Biology Provide Insights into Human Disease. Endocrine Reviews 22:2–35.

30. Sen, S., A. Dey, and U. Maulik, 2018. Identifying Potential Hubs for Kidney Renal Clear Cell Carcinoma from TF-miRNA-Gene Regulatory Networks. In Proceedings of IEEE Applied Signal Processing Conference 240–244.

31. Meszaros, B., G. Erdos, and Z. Dosztanyi, 2018. IUPred2A: context-dependent prediction of protein disorder as a function of redox state and protein binding. Nucleic Acids Research 46:W329–W337.

32. Xue, B., R. L. Dunbrack, R. W. Williams, A. K. Dunker, and V. N. Uversky, 2010. PONDR-FIT: A Meta-Predictor of Intrinsically Disordered Amino Acids. Biochimica et Biophysica Acta 1804:996–1010.

33. Zhang, Y., 2008. I-TASSER server for protein 3D structure prediction. BMC Bioinformatics 9.

34. Yang, J., R. Yan, D. X. A Roy, J. Poisson, and Y. Zhang, 2015. The I-TASSER Suite: Protein structure and function prediction. Nature Methods 12:7–8.

35. Croft, D., G. O’Kelly, and et al., 2011. Reactome: a database of reactions, pathways and biological processes. Nucleic Acids Research D691–D697.

36. Chen, C., S. Zabad, H. Liu, W. Wang, and C. Jeffery, 2018. MoonProt 2.0: an expansion and update of the moonlighting proteins database. Biochimica et Biophysica Acta 46:D640–D644.

37. Vazquez-Arreguín, K., C. Bensard, J. C. Schell, E. Swanson, X. Chen, J. Rutter, and D. Tantin, 2019. Oct1/Pou2f1 is selectively required for colon regeneration and regulates colon malignancy. PLoS Genetics 15:e1007687.

38. Bhoumik, A., S. Takahashi, W. Breitweiser, Y. Shiloh, N. Jones, and Z. Ronai, 2005. ATM-dependent phosphorylation of ATF2 is required for the DNA damage response. Molecular cell 18:577–587.

